# An inducible genetic model of chronic hypoxic signaling in cardiomyocytes precipitates severe cardiomyopathy and remodeling

**DOI:** 10.1101/2025.04.09.647295

**Authors:** Chelsea M. Phillips, Michael J. Zeitz, Eric H. Sapp, Karim M. Abouelenein, James W. Smyth

## Abstract

Cardiovascular disease remains the leading cause of death globally, underscoring the need for physiologically relevant models to investigate mechanisms of heart failure and arrhythmia. Chronic activation of hypoxic signaling pathways, particularly via the hypoxia-inducible factor (HIF) axis, is a key contributor to cardiac remodeling under stress. A major regulator of HIF signaling is the von Hippel-Lindau tumor suppressor (VHL) which, under normoxic conditions, targets HIFα for proteasomal degradation. Loss of VHL results in HIFα accumulation and persistent hypoxic signaling, but constitutive cardiomyocyte-specific *Vhl* knockout models are confounded by developmental effects and early mortality. Here, we develop and characterize an inducible, cardiomyocyte-specific *Vhl* knockout mouse model as a non-invasive and temporally controlled system to study chronic hypoxic stress and its contribution to cardiac remodeling and disease. *VhlLoxP/LoxP*;αMHC-MerCreMer^+/−^ mice were administered tamoxifen to induce *Vhl* deletion in adult cardiomyocytes. Within 5–7 days post-induction, mice displayed reduced ejection fraction, increased cardiac diameter, and elevated expression of cardiac stress markers. Transcriptomic and protein analyses revealed downregulation of key genes involved in cardiac structure and electrophysiology, including *Gja1* (Cx43), *Cdh2* (N-cadherin), *Cacna1c* (Ca_V_1.2), and *Kcnq1*. Importantly, these changes preceded overt cardiac remodeling, as confirmed in an abbreviated tamoxifen protocol. This inducible *Vhl* knockout model recapitulates hallmark features of dilated cardiomyopathy and highlights a subset of cardiac structural and ion channel genes as sensitive early responders to chronic hypoxic stress. This platform enables mechanistic dissection of disease onset and progression in ischemic heart disease and serves as a well-controlled and reproducible model for evaluating novel therapeutic strategies.

## Introduction

Cardiovascular disease is the leading cause of death in the United States and globally, with ischemic heart disease being the most common cause of heart failure^1, 2, 3, 4^. This underscores the importance of understanding molecular mechanisms driving cardiovascular disease pathogenesis and progression, as this will allow for the identification of novel substrates for therapeutic development. Elucidating the molecular underpinnings of cardiovascular disease requires the development of physiologically relevant animal models.

Encompassing a broad range of pathologies, cardiovascular disease is categorized as either ischemic or non-ischemic. Ischemic heart disease stems from a narrowing of coronary arteries resulting in reduced blood flow and oxygen to the heart. Resulting activation of hypoxic-stress related pathways in cardiomyocytes elicits pathological arrhythmogenic remodeling affecting transcription, translation, trafficking, function, and stability of critical ion channels and structural proteins^5–8^. Existing surgical mouse models have been used to investigate molecular mechanisms contributing to the pathologies of ischemic heart disease, with ligation of the left anterior descending artery (LAD) being the most common^9^. While this model reliably induces myocardial infarction, there are several limitations including, but not limited to, technical difficulty, variability in infarct size, and the confounding influence of the inflammatory response from surgery.

Aside from surgical approaches, genetic induction of cardiac stress is another method to model ischemic heart disease *in vivo*, with one of the most prominent approaches including genetic activation of the hypoxia inducible factor (HIF) pathway. Activity of the HIF basic-helix-loop-helix-PAS transcription factor, comprising an oxygen-sensitive α subunit and a constitutively expressed β subunit, results in the transcriptional activation of hypoxic stress response genes^10, 11^. During normoxic conditions, prolyl hydroxylase (PHD)-mediated hydroxylation of HIFα allows von Hippel-Lindau (VHL), a component of the E3 ubiquitin ligase complex, to ubiquitinate HIFα, leading to its downstream proteasomal degradation^12–16^. Hypoxia, however, renders PHDs enzymatically inactive^12^. This prevents hydroxylation and subsequent ubiquitination of HIFα and leads to its accumulation. HIFα can then translocate to the nucleus, where it dimerizes with HIFβ and binds to hypoxia-response elements (HRE) to modulate gene expression in response to the hypoxic cellular environment^17^.

Indeed, genetically manipulating the HIF pathway does recapitulate ischemic cardiovascular disease *in vivo*^18, 19^. Transcriptionally overexpressing HIF-1α in the myocardium reveals chronic HIF-1α signaling leads to cardiomyopathy, with abnormalities in capillary area, metabolism, and Ca^2+^ handling^18^. Primary regulation of HIFα occurring posttranslationally, however, suggesting deletion of upstream HIF pathway mediators to activate HIF signaling may be a more appropriate method for inducing cardiac stress. As such, manipulating HIF signaling via *Phd* deletion does mimic ischemic cardiomyopathy *in vivo*, but redundancy in the functions of the PHD paralogs (e.g., PHD1, 2, and 3) requires double knockout animals to achieve this phenotype^19^. A more straightforward method of inducing chronic stress signaling via pathway activation involves deletion of the tumor suppressor protein VHL. Mutations in *VHL* cause VHL syndrome, an autosomal dominant hereditary disease associated with tumor development, including pheochromocytomas^20^. While cardiac manifestations of VHL syndrome are rare, case studies demonstrate the presence of cardiac symptoms, including dilated cardiomyopathy, in a subset of VHL syndrome patients^21–23^. In some instances, surgical resection of the pheochromocytoma resolves the cardiac symptoms; however, cases where cardiac symptoms persist post-pheochromocytoma resection have been documented, suggesting chronic HIF signaling due to loss of functional VHL expression may underlie these cardiac manifestations of VHL syndrome^22, 23^.

Accordingly, mouse models of constitutive, cardiomyocyte-specific *Vhl* deletion result in cardiac dysfunction and increased mortality compared to wild-type controls^19, 24^. While this mouse model highlights the importance of VHL expression in cardiomyocytes and elucidates cardiomyocyte-specific VHL functions, using the constitutive mouse model to investigate mechanisms of ischemic cardiovascular disease proves challenging: nearly half of mice developed cardiac neoplasms capable of metastasis, making it difficult to disentangle the cardiac phenotypes and molecular signatures arising from cardiac neoplasms from those due to chronic hypoxic stress^24^. Moreover, the lack of temporal control coupled with early mortality introduces difficulty in dissecting mechanisms contributing to cardiovascular disease development from those involved in cardiovascular disease progression. Instead, an inducible, cardiomyocyte-specific *Vhl* knockout mouse model may prove to be a more favorable approach in inducing chronic cardiac hypoxic stress to non-invasively model cardiovascular disease. As such, we set out to generate an inducible, cardiomyocyte-specific *Vhl* knockout mouse model to create a non-invasive, reproducible mouse model of chronic cardiac hypoxic stress, which would allow mechanisms of cardiovascular disease pathogenesis and progression to be interrogated.

Here, we demonstrate that cardiomyocyte-specific *Vhl* deletion results in rapid cardiac remodeling comparable to dilated cardiomyopathy, as indicated by decreased ejection fraction and fractional shortening and increased cardiac diameter. Loss of VHL also results in dysregulated expression of key cardiac genes, an event preceding gross cardiomyopathy. Overall, we find that inducing chronic hypoxia in cardiomyocytes through *Vhl* deletion results in a non-invasive, reproducible mouse model of dilated cardiomyopathy, which ultimately allows for investigating molecular mechanisms contributing to dilated cardiomyopathy pathogenesis and progression.

## Results

### Cardiomyocyte-specific loss of VHL induces severe cardiomyopathy

To determine whether cardiomyocyte-specific *Vhl* deletion yields a viable, noninvasive mouse model of cardiomyopathy, we generated an inducible, conditional *Vhl* knockout (*Vhl*^−/−^) mouse line through crossing *Vhl-LoxP/LoxP* mice, containing loxP sites flanking the promoter and exon1 of *Vhl*, with an αMHC-MerCreMer^+/−^ mouse line, which expresses tamoxifen-inducible Cre recombinase under the control of the alpha-myosin heavy chain promoter. At 8 to 12 weeks, Cre activity, resulting in *Vhl* excision, was induced through daily administration of intraperitoneal (i.p.) tamoxifen (TMX) for five days (Figure 1A). Experimental controls included *Vhl-LoxP/LoxP*;αMHC-MerCreMer^+/−^ mice receiving daily vehicle injections (vehicle control), and *Vhl-LoxP/LoxP* mice receiving daily TMX injections (TMX control). Cardiac tissue was harvested at day 8 and used to confirm successful *Vhl* deletion. Protein levels of VHL were assessed through western blotting, which demonstrates reduced VHL protein expression in *Vhl*^−/−^ cardiac tissue compared to the controls (Figure 1B). Reductions in VHL expression are modest, likely due to its expression in the other cell types comprising cardiac tissue (e.g., cardiac fibroblasts and endothelial cells). To further confirm VHL loss, we investigated cardiac expression of hypoxia-inducible factor 2α (HIF-2α), a downstream target of VHL. As VHL targets HIFα for proteasomal degradation, loss of VHL expression results in HIFα accumulation in normoxic conditions. Western blotting demonstrates significant increases in HIF-2α expression in *Vhl*^−/−^ cardiac tissue compared to controls (Figure 1B). Together, these data confirm *Vhl* deletion and the subsequent onset of hypoxic signaling following TMX administration.

**Figure 1.**
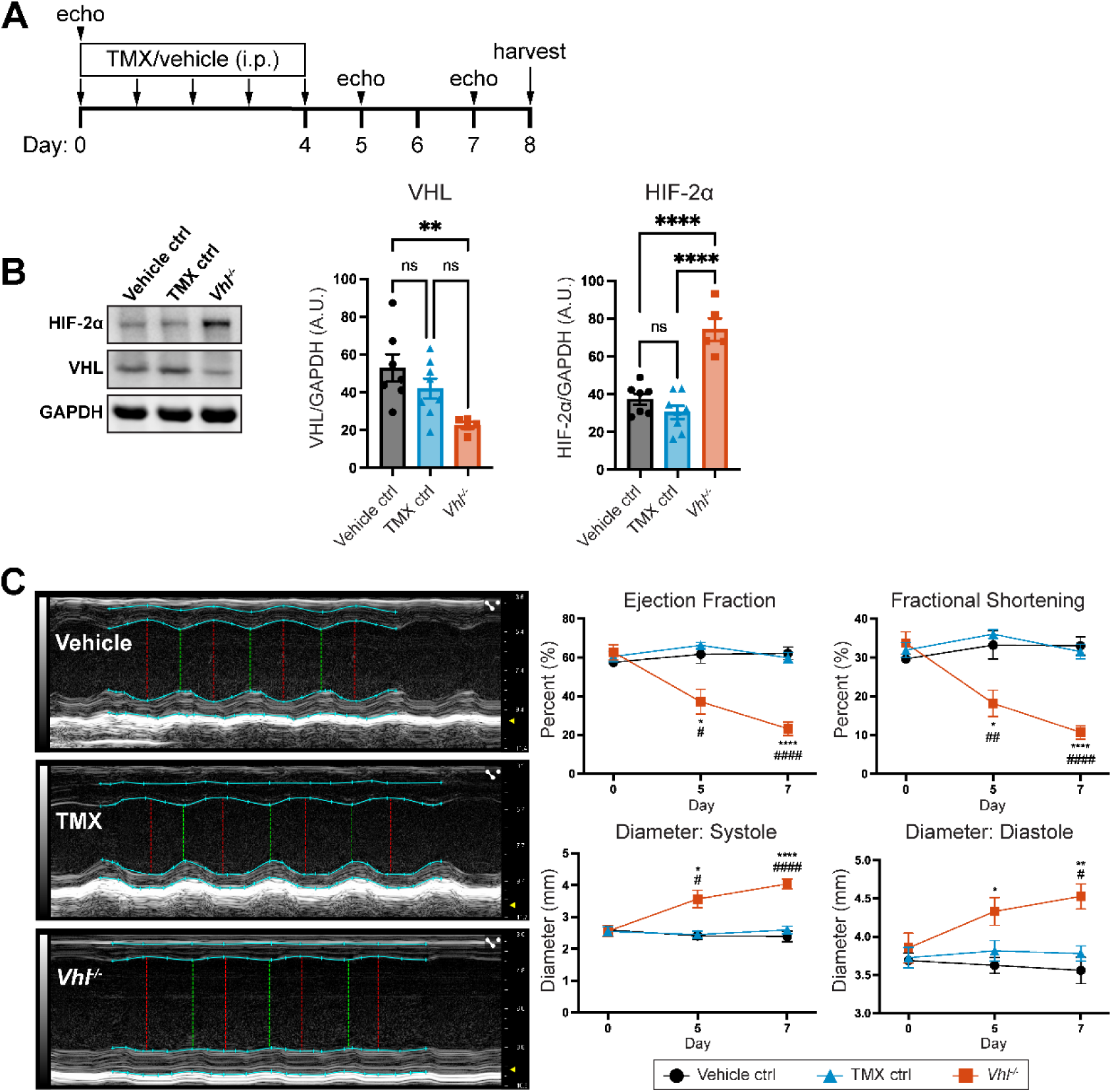
Cardiomyocyte-specific *Vhl* deletion induces severe cardiomyopathy. A) Schematic of experimental timeline. Mice underwent baseline echocardiography prior to receiving i.p. injections of TMX or vehicle for 5 days. Post-TMX administration, echocardiography was conducted at days 5 and 7, and cardiac tissue was harvested at day 8. B) Western blot of cardiac tissue isolated at day 8 for expression of VHL and HIF-2α, with GAPDH as a loading control (n = 7 vehicle control, 8 TMX control, and 5 *Vhl*^−/−^ samples). Quantification of VHL (left) and HIF-2α (right), with each normalized to GAPDH. One-way ANOVA with Tukey’s multiple comparisons test, \**P* < 0.05, \*\**P* < 0.01, and \*\*\*\**P* < 0.0001. C) Representative M-mode peristernal long axis (PSLAX) echocardiograms from day 7. Echocardiography quantifications include ejection fraction, fractional shortening, and cardiac diameter during systole and diastole (n = 7 vehicle control, 8 TMX control, and 6 *Vhl^−/−^* mice). Two-way repeated measures ANOVA with Geisser-Greenhouse correction, followed by Tukey’s multiple comparisons test. Compared to vehicle control: \**P* < 0.05 and \*\*\*\**P* < 0.0001. Compared to TMX control: #*P* < 0.05, ##*P* < 0.01, ####*P* < 0.0001.

To determine whether cardiomyocyte-specific *Vhl* deletion impacts cardiac structure and function, we conducted longitudinal echocardiography on days 0 (baseline, prior to TMX administration), 5, and 7 (Figure 1A). At baseline, ejection fraction, fractional shortening, and cardiac diameter are comparable across experimental groups (Figure 1C). At day 5, we observe alterations in cardiac function and structure in *Vhl*^−/−^ mice, as indicated by reduced ejection fraction and fractional shortening and increased cardiac diameter. These functional and structural cardiac abnormalities persist at day 7 (Figure 1C). Overall, the impaired cardiac function coupled with increased cardiac diameter suggests that cardiomyocyte-specific *Vhl* deletion results in a cardiac phenotype comparable to dilated cardiomyopathy.

### Loss of Cx43 expression occurs in the absence of VHL

With cardiomyocyte-specific *Vhl*^−/−^ deletion resulting in drastic cardiac remodeling, we next interrogated how loss of VHL affected expression of genes critical to cardiac structure and function using RT-qPCR. We first investigated transcript expression of the cardiac stress markers *Nppa* and *Nppb*, observing significant increases in mRNA levels of both cardiac stress markers in *Vhl*^−/−^ cardiac tissue compared to vehicle control and TMX control (Figure 2A). Moreover, we observe a switching of myosin heavy chain expression, as indicated by a significant decrease in *Myh6* transcript levels with a concomitant increase in *Myh7* transcript levels in *Vhl*^−/−^ cardiac tissue (Figure 2B)^25^. This observed switching of myosin heavy chain gene expression is consistent with observations in rodent models of cardiac hypertrophy^26, 27^.

**Figure 2.**
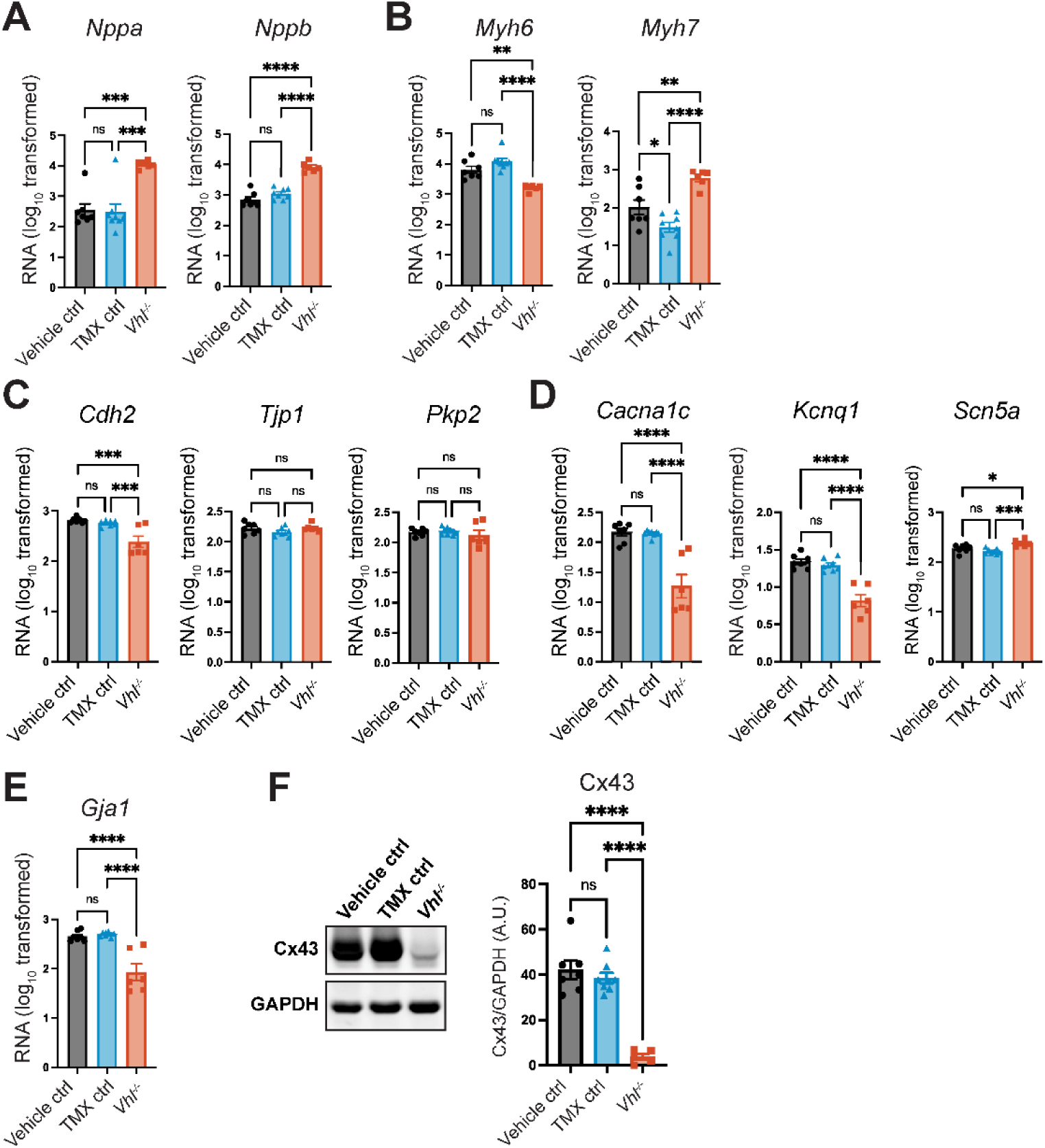
Loss of VHL expression alters expression of essential cardiac genes. RT-qPCR was employed to measure cardiac transcript expression of the following genes: A) cardiac stress markers *Nppa* and *Nppb*; B) myosin heavy chain genes *Myh6* and *Myh7*; C) intercalated disc junctional genes *Cdh2*, *Tjp1*, and *Pkp2*; D) cardiac ion channel genes *Cacna1c*, *Kcnq1*, and *Scn5a*; and E) gap junction gene *Gja1*. Transcript expression was normalized to *Gapdh*, with data representing log_10_ transformed values (n = 7 vehicle control, 8 TMX control, and 6 *Vhl^−/−^* samples). One-way ANOVA with Tukey’s multiple comparisons test, \**P* < 0.05, \*\**P* < 0.01, \*\*\**P* < 0.001, and \*\*\*\**P* < 0.0001. F) Western blot of cardiac tissue probing for expression of the gap junction protein Cx43. Quantification of Cx43 expression (right), normalized to GAPDH (n = 7 vehicle control, 8 TMX control, and 5 *Vhl^−/−^* samples). One-way ANOVA with Tukey’s multiple comparisons test, \*\*\*\**P* < 0.0001.

Due to the severe cardiac abnormalities in the *Vhl*^−/−^ mice, we also investigated transcript expression of key junctional genes required for intercalated disc structure. These include *Cdh2* (N-cadherin; adherens junctions), *Tjp1* (ZO-1; scaffolding protein), and *Pkp2* (Plakophilin 2; desmosome) (Figure 2C). Interestingly, transcript levels of *Cdh2* are reduced in *Vhl*^−/−^ cardiac tissue compared to control cardiac tissue, indicating disruption of the intercalated disc may be occurring following induction of chronic hypoxic stress due to *Vhl* deletion. Transcript levels of *Tjp1* and *Pkp2* remain unchanged in the *Vhl*^−/−^ cardiac tissue, suggesting targeted disruption of junctional gene expression does not arise from global transcriptional repression. In addition to the junctional genes profiled, we also investigated expression of genes encoding ion channels involved in cardiac electrophysiology, including *Cacna1c* (Ca_V_1.2), *Kcnq1* (KQT member 1, K_V_7.1), and *Scn5a* (Na_V_1.5) (Figure 2C). Strikingly, *Vhl*^−/−^ cardiac tissue demonstrates reduced transcript levels of *Cacna1c* and *Kcnq1* compared to controls; however, an increase in transcript expression of *Scn5a* is also observed. When investigating transcript expression of the gap junction gene *Gja1* (Connexin43; Cx43), which encodes the predominant connexin within the working myocardium^28^, we find significantly decreased *Gja1* transcript levels compared to vehicle and TMX controls (Figure 2E). Because of the significantly downregulated *Gja1* transcript levels in *Vhl^−/−^* cardiac tissue, we assessed Cx43 protein levels by western blotting. In accordance with the mRNA levels, Cx43 protein levels are significantly reduced in *Vhl*^−/−^ cardiac tissue compared to controls (Figure 2F). Together, these data demonstrate that chronic hypoxic signaling in cardiomyocytes decreases expression of genes crucial for cardiac structure and function. In the instance of Cx43, this decrease is observed at the protein level as well, likely affecting both action potential generation and propagation.

### Abbreviated TMX administration schedule deletes *Vhl* without cardiac remodeling

The decreased expression of key cardiac genes led us to ask whether these molecular changes preceded or followed the cardiac remodeling observed in the *Vhl*^−/−^ mice. To explore this, we first established a pre-cardiac remodeling timepoint through employing an abbreviated TMX administration schedule (Figure 3A). At day 0, echocardiography was conducted to establish baseline cardiac function and structure, followed by three days of TMX or vehicle i.p. injections. At day 3, echocardiography was conducted to determine whether *Vhl* deletion altered cardiac function or structure, and cardiac tissue was harvested. PCR genotyping of cardiac tissue at day 3 demonstrates *Vhl* was successfully deleted from cardiomyocytes (Figure 3B). Despite successful deletion of *Vhl* with this abbreviated timeline of TMX administration, echocardiography reveals cardiac functional and structural parameters remained unchanged in *Vhl^−/−^* mice compared to vehicle and TMX controls at day 3 (Figure 3C). Specifically, *Vhl*^−/−^ mice and control mice have comparable ejection fraction, fractional shortening, and cardiac diameter. Together, confirmation of *Vhl* deletion without altered cardiac structure and function allowed us to confidently use this abbreviated TMX administration schedule to interrogate molecular changes following cardiomyocyte-specific *Vhl* deletion prior to cardiac remodeling.

**Figure 3.**
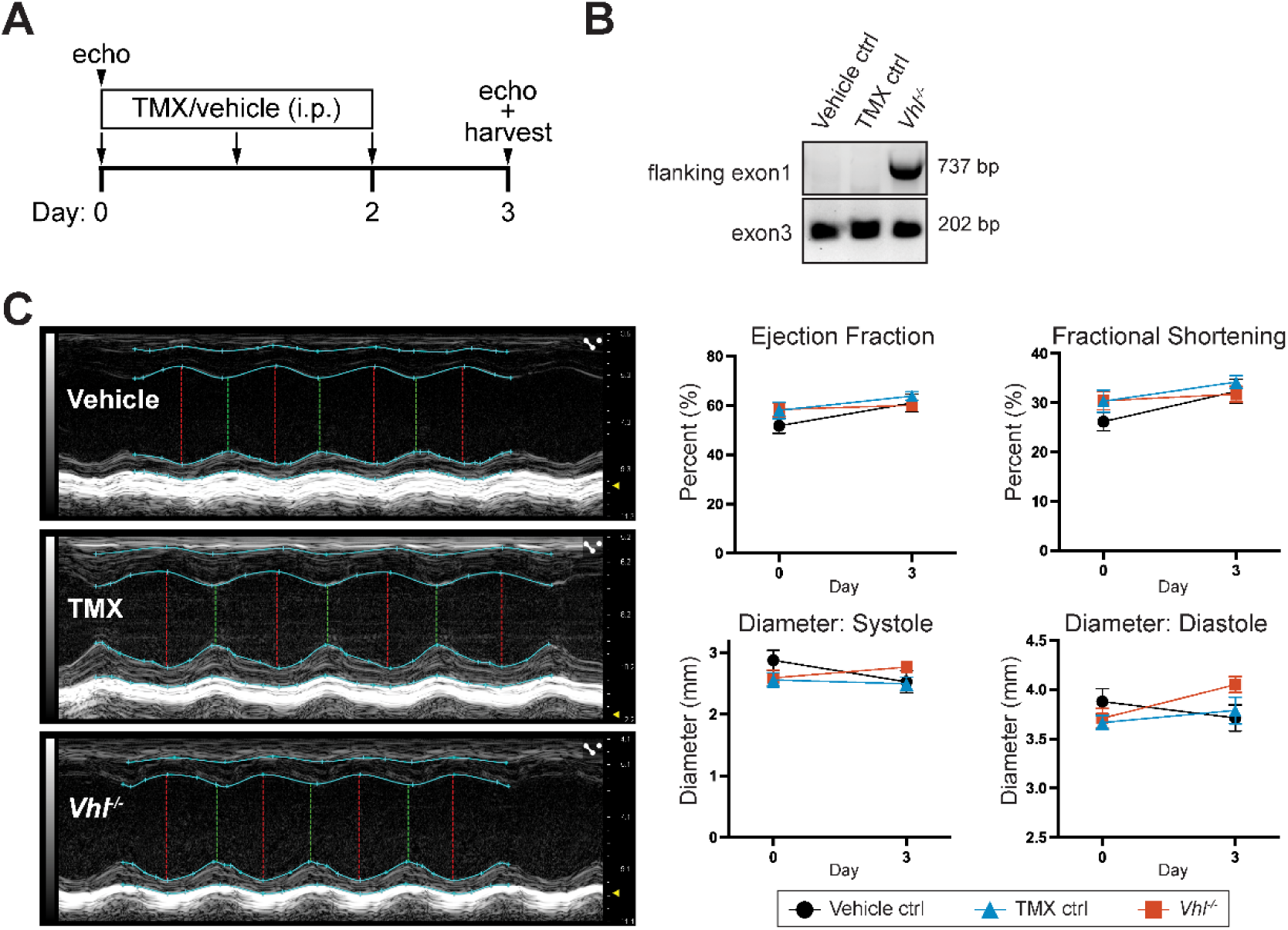
Abbreviated TMX administration results in VHL loss without cardiac remodeling. A) Schematic of experimental timeline. Mice underwent echocardiography prior to 3 days of daily i.p. TMX or vehicle administration. At day 3, echocardiography was conducted, and cardiac tissue was harvested. B) *Vhl* deletion was confirmed through genotyping cardiac tissue post-TMX administration. Representative DNA gel image demonstrates the presence of a deletion band in *Vhl*^−/−^ cardiac tissue, with exon3 as a genotyping control C) Representative M-mode PSLAX echocardiograms recorded at day 3, with quantifications including ejection fraction, fractional shortening, and cardiac diameter during systole and diastole (n = 6 vehicle control, 8 TMX control, and 7 *Vhl^−/−^* mice). Two-way repeated measures ANOVA with Geisser-Greenhouse correction, followed by Tukey’s multiple comparisons test.

### Cardiac transcriptomic alterations precede gross cardiomyopathy in *Vhl*^−/−^ hearts

To investigate the molecular changes upon VHL loss prior to gross cardiomyopathy, we conducted RNA-sequencing (RNA-seq) on RNA isolated from *Vhl*^−/−^ and TMX control cardiac tissue isolated at day 3. Compared to TMX control, 816 genes are differentially expressed in the *Vhl*^−/−^ cardiac tissue (adjusted p-value ≤ 0.05 and absolute value of log2 fold change ≥ 1.0) (Figure 4A). Of these 816 differentially expressed genes (DEGs), 398 are upregulated (48.8%), and 418 are downregulated (51.2%). Overall, this indicates that while key cardiac genes are downregulated in the absence of VHL, this is not due to global transcriptional repression. To understand how loss of VHL alters the cardiac transcriptomic landscape prior to cardiac remodeling, we first explored the most significant DEGs, as indicated by adjusted p-value (Table 1). Strikingly, this includes the cardiac intercalated disc genes *Cdh2* and *Gja1*, both of which are downregulated in *Vhl^−/−^* cardiac tissue compared to TMX control tissue. Furthermore, other genes within the top 20 most significant DEGs include those involved in calcium handling, such as *Cacna1c*, *Ryr2*, and *Trdn* (encoding a RYR2-interacting integral membrane protein), all of which are downregulated. Aside from altered expression of genes required for calcium handling, *Vhl*^−/−^ cardiac tissue also demonstrates downregulation of the ion channel *Slc8a1*. *Fgf13*, demonstrated to be involved in Na_V_1.5 regulation, is also downregulated in *Vhl*^−/−^ hearts^29^. While transcript expression of genes essential for proper cardiac function is downregulated following VHL loss, a subset of genes within the top 20 most significant DEGs are associated with cardiovascular disease. These include *Fhod3*, mutations in which are associated with dilated cardiomyopathy, along with *Alkp3*, *Ccn2*, and *Tnfrsf12a*, all of which are implicated in cardiac hypertrophy^30–33^. Likewise, we see downregulation of *Tecrl*, implicated in inherited cardiac arrhythmias^34^. We also observe upregulation of the oxidative stress response gene *Srxn1*, which is unsurprising given *Vhl* deletion results in chronic hypoxic stress signaling.

**Figure 4.**
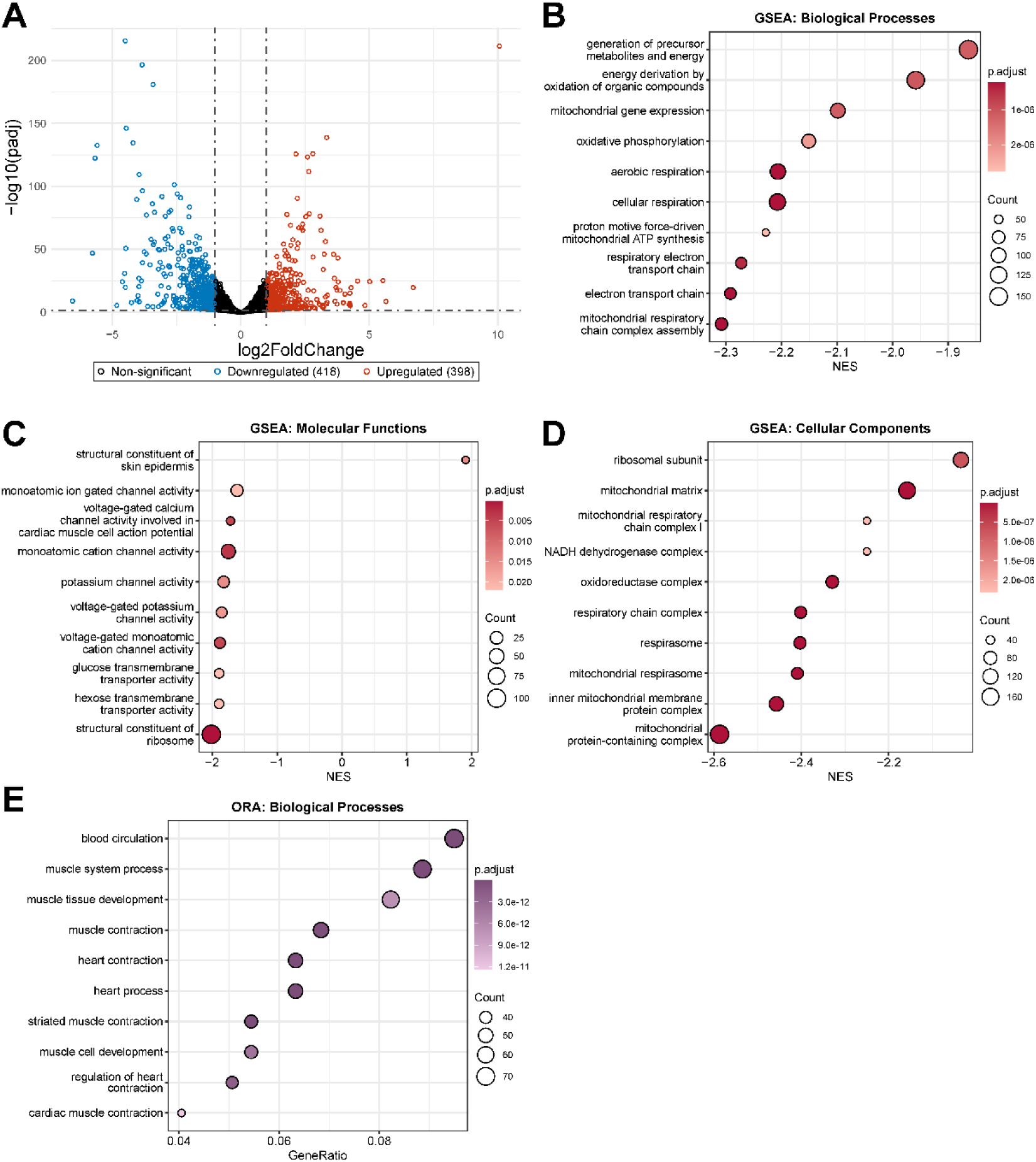
Cardiomyocyte-specific *Vhl* deletion suppresses expression of genes essential for cardiac function. A) RNA-seq was conducted to identify differentially expressed genes (DEGs) in cardiac tissue isolated at day 3 from *Vhl*^−/−^ mice compared to TMX control. Data are represented as a volcano plot. Significantly upregulated DEGs are highlighted in red, while significantly downregulated DEGs are highlighted in blue (n = 3). Significant DEGs: adjusted p-value (padj) ≤ 0.05 and |log2FoldChange| ≥ 1.0. Gene set enrichment analysis (GSEA) was conducted to identify enriched GSEA terms. The top ten enriched GSEA terms are represented as dot plots for B) biological processes, C) molecular functions, and D) cellular components. GSEA terms are arranged by normalized enrichment score (NES), with dot size indicating the number of genes and color representing padj. GSEA terms were identified as enriched if padj ≤ 0.05. E) Over-representation analysis (ORA) was conducted with the significant DEGs to identify enriched biological processes gene ontology (GO) terms. The top ten enriched GO terms are represented as a dot plot. GO terms are arranged by gene ratio, with dot size indicating gene count and padj value indicated by color. GO terms are considered enriched if padj ≤ 0.05.

**Table 1.**
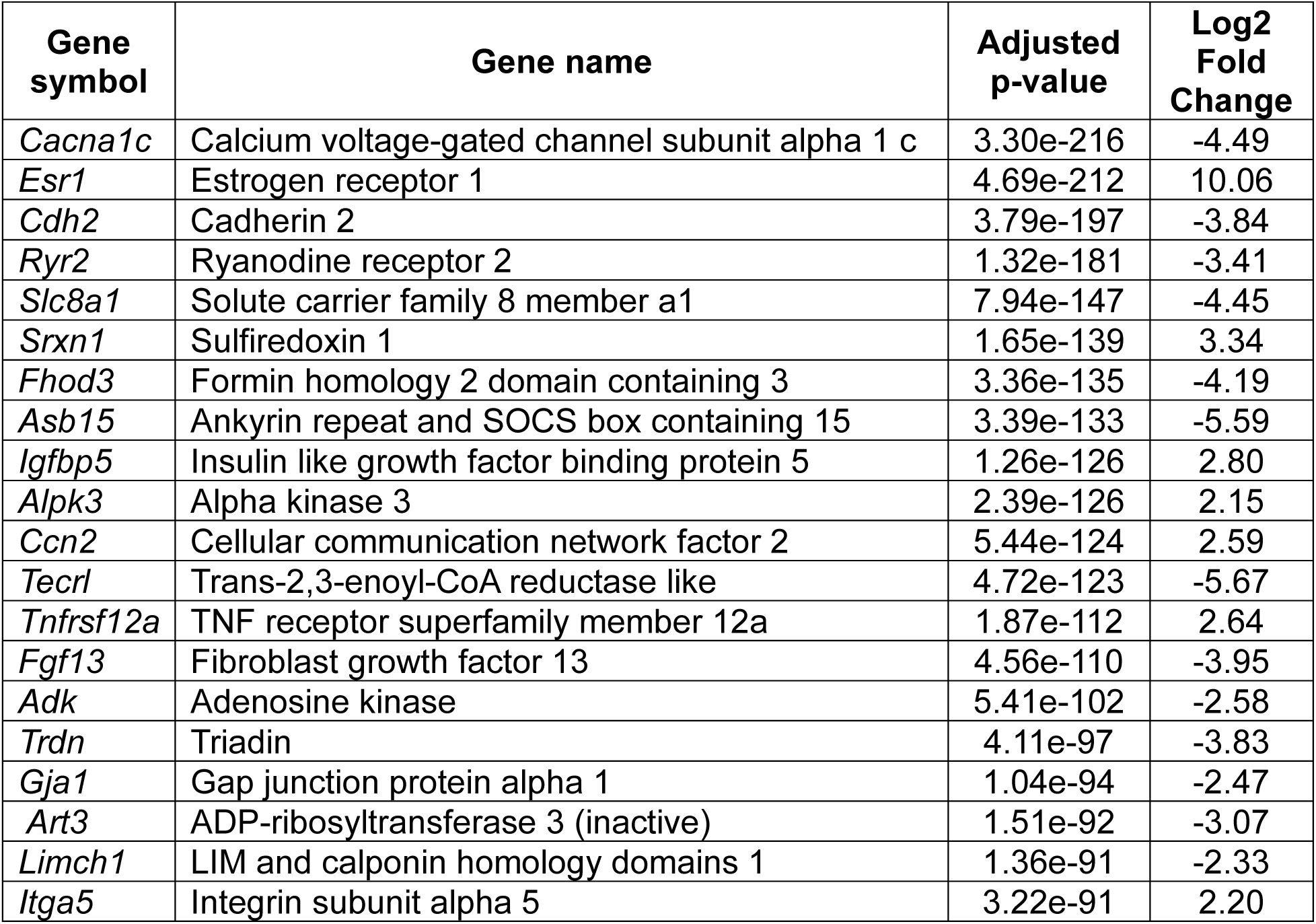
Most significant differentially expressed genes in *Vhl*^−/−^ cardiac tissue compared to TMX control.

| Gene symbol | Gene name | Adjusted p-value | Log2 Fold Change |
| --- | --- | --- | --- |
| *Cacna1c* | Calcium voltage-gated channel subunit alpha 1 c | 3.30e-216 | -4.49 |
| *Esr1* | Estrogen receptor 1 | 4.69e-212 | 10.06 |
| *Cdh2* | Cadherin 2 | 3.79e-197 | -3.84 |
| *Ryr2* | Ryanodine receptor 2 | 1.32e-181 | -3.41 |
| *Slc8a1* | Solute carrier family 8 member a1 | 7.94e-147 | -4.45 |
| *Srxn1* | Sulfiredoxin 1 | 1.65e-139 | 3.34 |
| *Fhod3* | Formin homology 2 domain containing 3 | 3.36e-135 | -4.19 |
| *Asb15* | Ankyrin repeat and SOCS box containing 15 | 3.39e-133 | -5.59 |
| *Igfbp5* | Insulin like growth factor binding protein 5 | 1.26e-126 | 2.80 |
| *Alpk3* | Alpha kinase 3 | 2.39e-126 | 2.15 |
| *Ccn2* | Cellular communication network factor 2 | 5.44e-124 | 2.59 |
| *Tecrl* | Trans-2,3-enoyl-CoA reductase like | 4.72e-123 | -5.67 |
| *Tnfrsf12a* | TNF receptor superfamily member 12a | 1.87e-112 | 2.64 |
| *Fgf13* | Fibroblast growth factor 13 | 4.56e-110 | -3.95 |
| *Adk* | Adenosine kinase | 5.41e-102 | -2.58 |
| *Trdn* | Triadin | 4.11e-97 | -3.83 |
| *Gja1* | Gap junction protein alpha 1 | 1.04e-94 | -2.47 |
| *Art3* | ADP-ribosyltransferase 3 (inactive) | 1.51e-92 | -3.07 |
| *Limch1* | LIM and calponin homology domains 1 | 1.36e-91 | -2.33 |
| *Itga5* | Integrin subunit alpha 5 | 3.22e-91 | 2.20 |

We also investigated transcriptomic changes through gene set enrichment analysis (GSEA), providing an overview of collective changes in gene expression (Figures 4B-D). GSEA reveals a striking downregulation of metabolic genes. The most significantly enriched biological process GSEA terms all relate to metabolism, such as generation of precursor metabolites and energy (p.adjust = 1.16e-06, NES = −1.86), energy derivation by oxidation of organic compounds (p.adjust = 1.07e-06, NES = −1.96), mitochondrial gene expression (p.adjust = 1.16e-06, NES = −2.10), oxidative phosphorylation (p.adjust = 2.18e-06, NES = −2.15), and respiratory electron transport chain (p.adjust = 4.45e-07, NES = −2.27) (Figure 4B). A negative enrichment score for metabolic terms was also observed for the most significant cellular components GSEA terms; examples include mitochondrial matrix (p.adjust = 1.56e-08, NES = −2.16), NADH dehydrogenase complex (p.adjust = 2.31e-06, NES = −2.25), oxidoreductase complex (p.adjust = 1.56e-08, NES = −2.33), and the respirasome (p.adjust = 1.56e-08, NES = −2.40) (Figure 4D). Significantly enriched molecular function GSEA terms, however, demonstrate negative enrichment of ion channel-related terms, such as monoatomic ion gated channel activity (p.adjust = 2.16e-02, NES = −1.62), voltage-gated calcium channel activity involved in cardiac muscle cell action potential (p.adjust = 5.10e-03, NES = −1.72), and potassium channel activity (p.adjust = 1.50e-02, NES = −1.82) (Figure 4C). Moreover, over-representation analysis (ORA) for biological process gene ontology (GO) terms demonstrates an enrichment of GO terms relating to cardiac function: blood circulation (p.adjust = 1.55e-15), heart contraction (p.adjust = 6.14e-16), heart process (p.adjust = 2.12e-15), and cardiac muscle contraction (p.adjust = 1.24e-11) (Figure 4E). Together, RNA-seq reveals that transcriptomic changes to key cardiac genes, including those involved in metabolism, ion channel activity, and cardiac function, occur prior to chronic hypoxic stress-induced cardiac remodeling in *Vhl*^−/−^ hearts.

### VHL loss alters transcript expression of cardiac ion channels

To further investigate the transcriptomic changes following loss of VHL expression within cardiomyocytes, we conducted an over-representation analysis for molecular function GO terms. Largely, we find significant enrichment of GO terms related to ion channel activity, such as monoatomic ion gated channel activity (p.adjust = 7.81e-03), suggesting altered transcript expression of ion channels required for proper cardiomyocyte function (Figure 5A). Within the *Vhl*^−/−^ hearts, 29 significant DEGs comprise monoatomic ion gated channel activity, with 12 DEGs upregulated and 17 downregulated (Figure 5B). A subset of the significant DEGs in this enriched GO term encode voltage-gated calcium channel subunits and their auxiliary subunits (Figure 5C). Examples of significant DEGs encoding voltage-gated calcium channel subunits include *Cacna1c* and *Cacna1s*, both of which are downregulated, and *Cacna1h*, which is upregulated in *Vhl*^−/−^ hearts compared to TMX control hearts. The same pattern is observed with the auxiliary subunits: *Cacna2d1* and *Cacnb2* are both downregulated in *Vhl*^−/−^ cardiac tissue, while *Cacna2d3* and *Cacng6* are both upregulated. The other major subset of genes comprising monoatomic ion gated channel activity are potassium channels (Figure 5D). Transcript expression of the predominant cardiac potassium channels is reduced in *Vhl*^−/−^ hearts compared to TMX controls; this is observed for *Kcnd2*, *Kcnj2*, *Kcnj3*, and *Kcnq1.* Collectively, this reveals that chronic hypoxic stress due to VHL loss alters expression of ion channels essential for cardiac electrophysiology.

**Figure 5.**
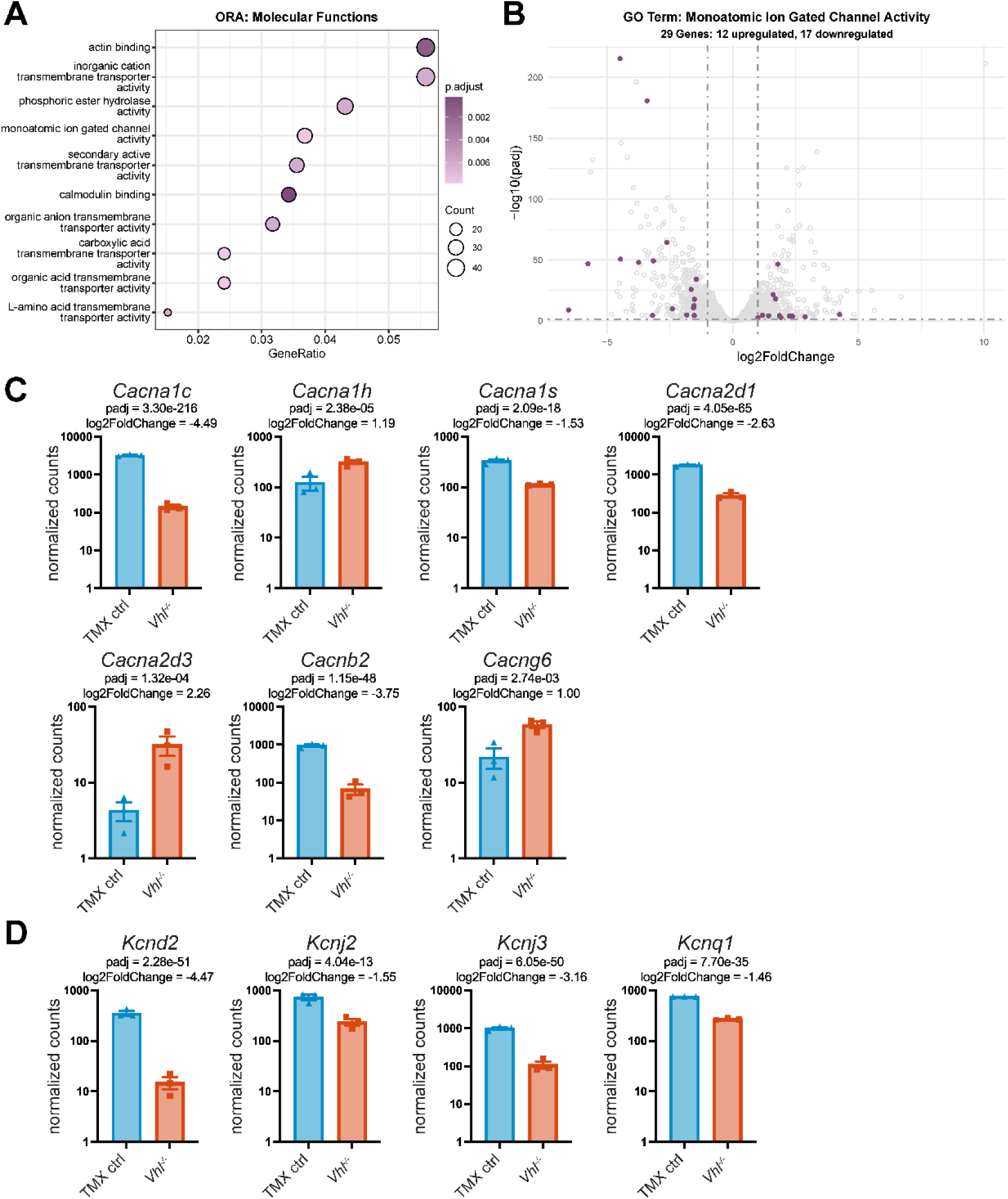
Cardiomyocyte-specific *Vhl* deletion alters expression of genes essential for normal cardiac electrophysiology. A) ORA was conducted to identify significantly enriched molecular function GO terms. Dot plot represents the top ten enriched GO terms arranged by gene ratio, with dot size and color indicating gene count and padj value, respectively. Molecular function GO terms were determined as significant if padj ≤ 0.05. B) Volcano plot representing all DEGs (as in Figure 4A); DEGs within the GO term monoatomic ion gated channel activity are highlighted in purple. Normalized RNA-seq gene counts were plotted for monoatomic ion gated channel activity genes of interest, including C) voltage-gated calcium channel subunits and D) potassium channels. DESeq2 statistics (i.e., padj and log2FoldChange) are included above each graph.

### *Vhl*^−/−^ hearts demonstrate downregulation of intercalated disc genes

When conducting an over-representation analysis for cellular components, we observe GO terms related to cardiomyocyte structures are significantly enriched (Figure 6A). Of interest is the significant enrichment of the GO term intercalated disc (p.adjust = 2.68e-05). Of the 15 significant DEGs within the GO term interacalated disc, 13 are downregulated in *Vhl*^−/−^ cardiac tissue compared to TMX control hearts (Figure 6B). These DEGs include junctional genes, such as *Cdh2* (N-cadherin) and *Gja1* (Cx43), both of which are downregulated in *Vhl*^−/−^ hearts (Figure 6C). Likewise, we observe downregulation of genes involved in the regulation of sodium channel expression, localization, and function, such as *Fgf13* and *Rangrf* (Figure 6D). Genes encoding proteins responsible for linking junctional proteins to the cytoskeleton also comprise this enriched GO term, with examples including *Ank2*, *Ctnna3*, *Dst*, and *Pkp2* (Figure 6E). Finally, we observe downregulation of *Tmem65*, a gene implicated in both gap junction and sodium channel function within cardiac tissue (Figure 6F)^35, 36^. Ultimately, investigating genes within the enriched GO term intercalated disc reveals that intercalated disc genes are primarily downregulated in *Vhl*^−/−^ hearts prior to gross cardiomyopathy.

**Figure 6.**
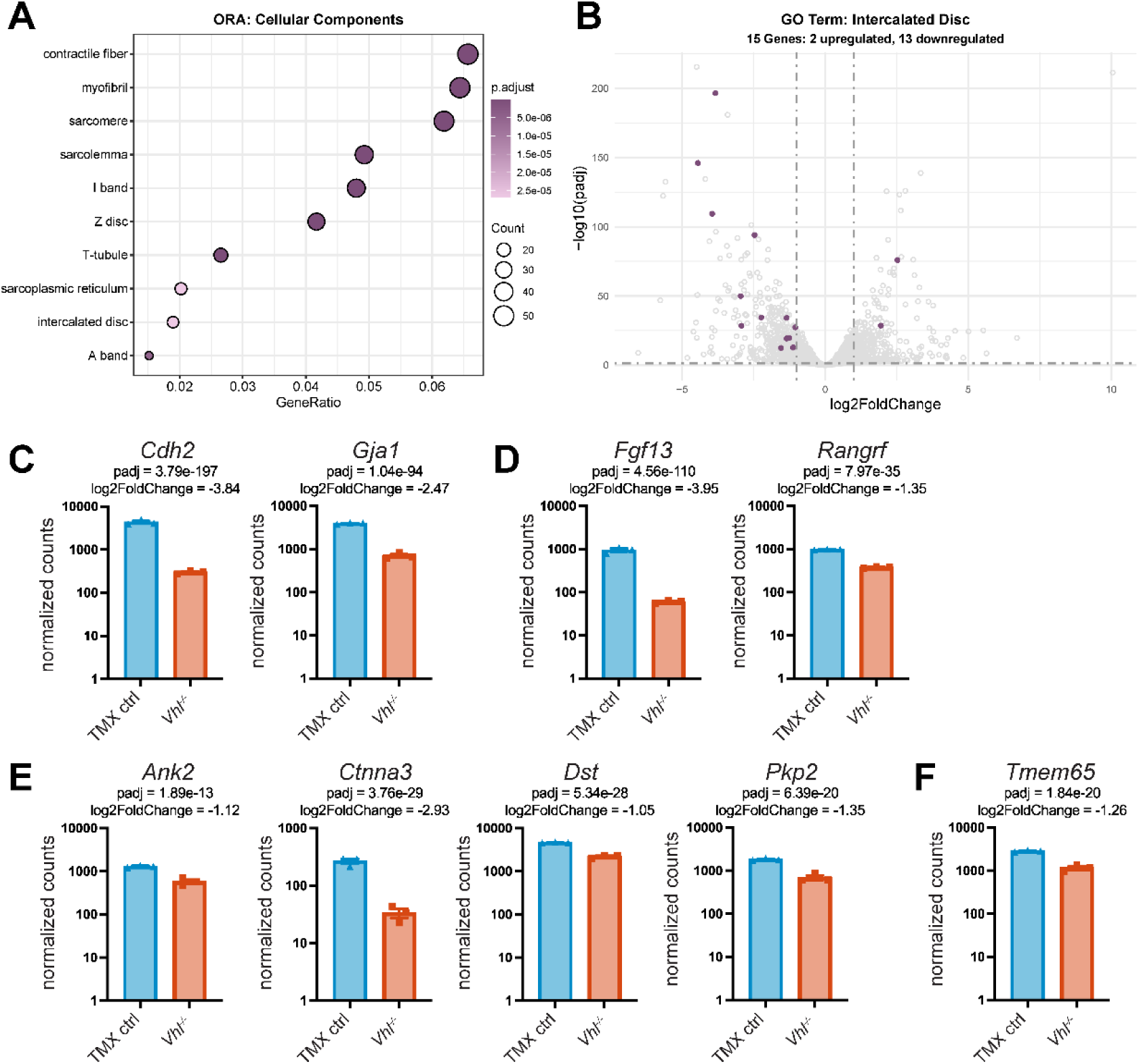
*Vhl* loss alters expression of genes encoding key cardiac structural genes. A) To determine significantly enriched cellular component GO terms, ORA was conducted. Dot plot demonstrates the top ten enriched GO terms arranged by gene ratio; dot size and color represent gene count and padj value, respectively. GO terms are classified as enriched if padj value ≤ 0.05. B) Volcano plot depicting differentially expressed genes (as in figure 4A), with intercalated disc DEGs represented in purple. Normalized RNA-seq gene counts were plotted for intercalated disc genes of interest, such as C) junctional genes, D) genes involved in sodium channel regulation, E) cytoskeletal linking genes, and F) *Tmem65*. DESeq2 statistics (i.e., padj and log2FoldChange) are included above each graph.

### Cx43 downregulation precedes cardiac remodeling in *Vhl*^−/−^ hearts

With RNA-seq demonstrating that the transcriptomic landscape of *Vhl*^−/−^ cardiac tissue is altered prior to gross cardiac remodeling, we sought to further characterize the molecular signatures of the *Vhl*^−/−^ cardiac tissue at this pre-remodeling time point, as these molecular changes may drive the cardiac remodeling. Through RT-qPCR, we find loss of cardiomyocyte VHL expression results in upregulation of the cardiac stress markers *Nppa* and *Nppb* without inducing a switch in the dominant myosin heavy chain gene from *Myh6* to *Myh7* (Figure 7A-B). This further confirms that the absence of VHL promotes cardiac stress, which arises prior to molecular remodeling of the cardiomyocyte structure. Additionally, we employed RT-qPCR to confirm our RNA-seq findings: we observe a significant reduction in *Gja1* transcript expression in *Vhl*^−/−^ cardiac tissue compared to vehicle and TMX controls (Figure 7C). Moreover, this translates to the protein level: *Vhl*^−/−^ mice have significantly decreased cardiac levels of Cx43 compared to the controls (Figure 7D). Together, this demonstrates that the decreased Cx43 mRNA and protein levels precedes cardiac structural and functional abnormalities observed in the *Vhl*^−/−^ mice, indicating that the reduction in Cx43 expression may be a key contributor to the gross cardiomyopathy observed in the *Vhl*^−/−^ mice.

**Figure 7.**
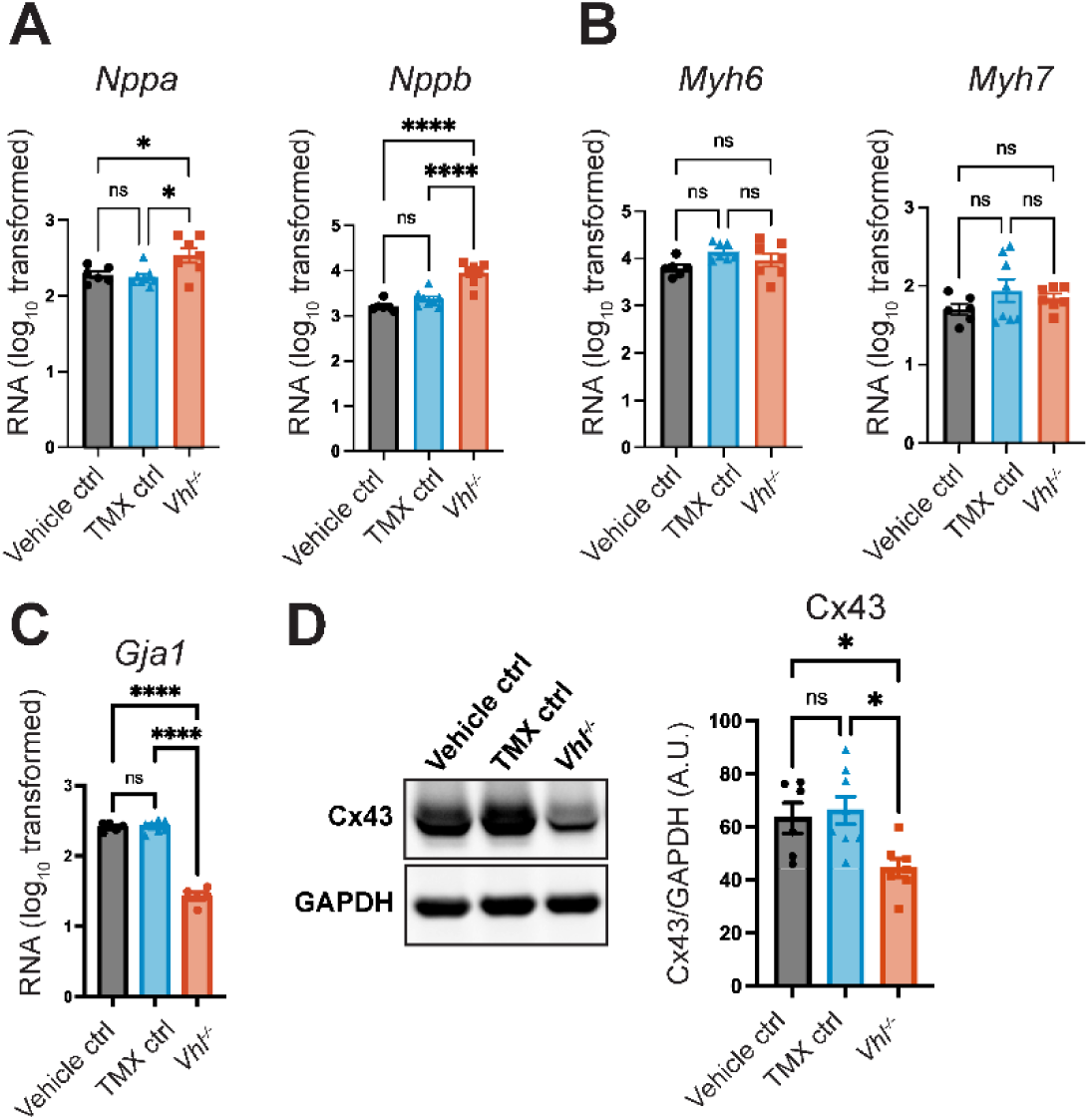
Cx43 downregulation precedes cardiac remodeling. RT-qPCR was employed to measure transcript expression of A) the cardiac stress markers *Nppa* and *Nppb*, B) the myosin heavy chain genes *Myh6* and *Myh7*, and C) the gap junction gene *Gja1*. Transcript expression was normalized to *Gapdh*. Data represent log_10_ transformed values. For A and B: n = 6 vehicle control, 7 TMX control, and 7 *Vhl^−/−^* samples. For C: n = 5 vehicle control, 7 TMX control, and 5 *Vhl*^−/−^ samples. One-way ANOVA with Tukey’s multiple comparisons test, \**P* < 0.05 and \*\*\*\**P* < 0.0001. D) Western blot probing for Cx43 expression in cardiac tissue isolated at day 3. Quantification of Cx43 expression normalized to GAPDH. (n = 6 vehicle control, 8 TMX control, and 7 *Vhl^−/−^* samples). One-way ANOVA with Tukey’s multiple comparisons test, \**P* < 0.05.

## Discussion

Through generation of an inducible, cardiomyocyte-specific *Vhl* knockout mouse model, we demonstrate that chronic hypoxic signaling within cardiomyocytes results in rapid cardiac remodeling analogous to that of dilated cardiomyopathy, as revealed by decreased cardiac function and increased cardiac diameter. Remarkably, genes central to cardiac structure and electrophysiology appear to be particularly susceptible to hypoxic stress signaling. Severe reductions in cardiac Cx43 transcript and protein levels, along other key cardiac genes, demonstrate that even isolated activation of hypoxic-signal transduction alone in cardiomyocytes is sufficient to essentially disassemble cardiac function at the cellular level. Chronic hypoxia-induced alterations to the cardiac transcriptomic landscape and downregulation of Cx43 protein levels, however, precede cardiac structural changes, suggesting these molecular signatures drive the cardiac remodeling. Together, our data highlight the inducible, cardiomyocyte-specific *Vhl* knockout mouse as a model for investigating molecular mechanisms driving cardiovascular disease development and progression, particularly within the context of ischemic cardiovascular disease.

The gap junction protein connexin43 (Cx43, gene name *Gja1*), is the predominant connexin expressed in the working myocardium^28^. Cx43 is crucial for cardiac action potential propagation and coordinated cardiac contraction; loss of Cx43 expression and its dispersion from the intercalated disc are observed across various cardiovascular diseases^37–40^. We find significantly reduced *Gja1* transcript and protein levels in *Vhl*^−/−^ cardiac tissue compared to vehicle and TMX controls prior to structural remodeling implicating its role in early-stage ischemic heart disease. While such losses in Cx43 expression, and therefore gap junction function, contribute to arrhythmia, it is likely such mechanisms evolved to prevent the spread of toxic metabolites and protect cardiac tissue during ischemic events.

While rare, case studies have reported incidences of dilated cardiomyopathy in individuals with VHL syndrome, indicating a potential link between expression of functional VHL and cardiac structure and function^21^. These studies have attributed these cardiac manifestations of VHL syndrome to pheochromocytomas, which are commonly associated with VHL syndrome. Indeed, surgical resection of the pheochromocytomas can be sufficient to restore cardiac function; however, instances where pheochromocytoma resection is insufficient to reverse the observed dilated cardiomyopathy phenotype have been described^22, 23^. VHL syndrome is a heterozygous condition, and in most cases may preclude high levels of cardiac HIF activity compared with homozygous VHL knockout. Furthermore, VHL syndrome-causing mutations, while all localized to the VHL gene, can result in different levels of HIF deregulation^41^. These factors may reduce the likelihood of presenting with a pathological cardiac phenotype that is so apparent in our data.

We have found that induced cardiomyocyte-specific loss of VHL expression directly underlies pathological cardiac remodeling. In accordance with our findings, cardiomyocyte-specific deletion of PHD2 and PHD3, enzymes responsible for HIF hydroxylation, leads to ischemic cardiomyopathy, marked by a thinning of the ventricle walls and increased ventricular diameter. Moreover, this particular study also demonstrated a similar, yet more severe, cardiac phenotype with cardiomyocyte-specific VHL deletion^19^. In another cardiomyocyte-specific VHL mouse model, it has been observed that concurrent deletion of HIF-1α prevents the development of cardiovascular complications/remodeling, demonstrating that these cardiac pathologies were not due to HIF independent effects of VHL^24^. Furthermore, in a mouse model of myocardial HIF-1α overexpression, it has been demonstrated that chronic HIF-1α upregulation, in combination with aging or increased mechanical load, leads to cardiomyopathy with abnormalities in capillary area, metabolism, and Ca^2+^ handling^18^. Interestingly, induced expression of a mutant version of HIF-1α resistant to hydroxylation in mouse hearts results in rapid ventricular dysfunction, illustrating the importance of stabilization of HIF-1α activity in generation of a severe pathological phenotype^42^.

Overall, this model system offers several advantages for investigating mechanisms of ischemic heart disease over existing models. Knocking out *Vhl* allows for the interrogation of downstream effects from all HIF isoforms expressed in the heart, rather than limiting the analysis to HIF-1α alone, as is typical in overexpression systems. This is especially valuable given the distinct yet sometimes overlapping roles of HIF-1α and HIF-2α in cardiovascular physiology and pathophysiology. Additionally, the inducible nature of this model provides precise temporal control over gene deletion, avoiding developmental confounders that often complicate interpretation in constitutive knockout models. This is particularly important for studying ischemic heart disease, which predominantly affects aging populations. Our approach allows the modeling of chronic hypoxic signaling at defined adult or aged time points, more accurately reflecting the clinical trajectory of human disease. Furthermore, the ability to isolate early molecular changes before the onset of overt remodeling offers a critical window for identifying candidate biomarkers or therapeutic targets. By recapitulating both the transcriptional and phenotypic hallmarks of dilated cardiomyopathy in a non-invasive and reproducible manner, this platform provides a powerful tool for dissecting the temporal progression of ischemic injury and its downstream remodeling processes. As such, it may inform mechanistic understanding of hypoxia-driven cardiac pathology and serve as a versatile foundation for preclinical evaluation of targeted therapies aimed at halting or reversing disease progression.

## Materials and Methods

### Animals

*Vhl-LoxP/LoxP* B6.129S4(C)-*Vhl^tm1Jae^*/J mice (Jackson Laboratories) were crossed with αMHC-MerCreMer^+/−^ mice to generate inducible cardiomyocyte-specific *Vhl* knockout (*Vhl*^−/−^) mice upon tamoxifen (75 mg/kg) treatment^43^.

### *Vhl* knockout genotyping

Heart tissue was homogenized using a Bead Mill 4 (Thermo Fisher) and DNA purified using DNeasy Blood and Tissue Kit (Qiagen) according to manufacturer’s instructions. Primers flanking the *loxP* sites surrounding the *Vhl* promoter and exon1 were used to detect *Vhl* deletion yielding a 737 bp product post-tamoxifen treatment. Positive control PCR primers targeting the unedited *Vhl* exon 3 produce a 202 bp product.

### Echocardiography

Echocardiography was performed with a Fuji Film Vevo F2 Imaging System with a UHF46x transducer. For experiments investigating cardiac remodeling, echocardiography was conducted at days 0, 5, and 7. For experiments investigating molecular changes prior to cardiac remodeling, echocardiography was performed at days 0 and 3. Mice were anesthetized in 3 % isoflurane in 2.5 % supplemental O_2_. Peristernal long axis (PSLAX) B-mode and M-mode images were acquired. Concurrently, temperature was monitored via rectal thermometer and cardiac electrical activity was obtained via surface electrodes. Data were analyzed in Vevo LAB (SW version 5.7.1) over 3.5 cardiac cycles, measuring ejection fraction (EF %), fractional shortening (FS %), and cardiac diameter during systole and diastole.

### RNA-sequencing

RNA was isolated from snap frozen cardiac ventricular tissue using TriZol. Tissue was homogenized in TRIzol (Thermo Fisher) using a bead mill 4 homogenizer (Thermo Fisher) followed by phenol-chloroform extraction. RNA was further purified with the PureLink RNA Mini Kit (Thermo Fisher) and residual DNA was removed by on column DNaseI digestion according to manufacturer recommendations. Library preparation, RNA-sequencing, and initial bioinformatics data analysis (e.g., data filtering, alignment, and generation of read counts matrices) were conducted by Innomics Inc. Briefly, sequencing libraries were generated as follows: after poly(A) enrichment, cDNA was synthesized via mRNA fragmentation and reverse transcription with random N6 primers. Following end-repair, 5’-phosphorylation, and adapter ligation, PCR amplification was conducted, with the resulting products denatured to form a single-stranded circular DNA library via bridged primer. Paired-end sequencing (read length 150 bp) was conducted on a DNBSEQ platform. Raw sequencing reads were filtered using SOAPnuke (v1.5.6) to remove reads with adapter contamination, a high percentage of unknown bases (greater than 0.1%), or low quality^44^. HISAT (v2.2.1) and Bowtie2 (v2.4.5) were used to align clean reads to mm39 *Mus musculus* genome (UCSC) and the mm39 *Mus musculus* gene set, respectively^45, 46^. Gene counts were obtained using RSEM (v1.3.1), with differentially expressed genes (DEGs) identified using the DESeq2 package (v1.42.1) in R (v4.3.1)^47, 48^. Log2 fold change (log2FC) shrinkage was conducted via the Ashr method (v2.2-63)^49^. Significant DEGs were defined as those with an adjusted p-value ≤ 0.05 and an absolute value of log2FC ≥ 1.0. Gene set enrichment (GSEA) and over-representation analysis (ORA) were both conducted with the R package clusterProfiler (v4.10.1)^50^. Briefly, GSEA was conducted with all DEGs, with p-values adjusted via the Benjamini-Hochberg method. ORA was conducted using significant DEGs (adjusted p-value ≤ 0.05 and absolute value of log2FC ≥ 1.0); p-values were adjusted using the Benjamini-Hochberg method. Data visualization was conducted using ggplot2 (v3.5.1) and DOSE (v3.28.2)^51, 52^.

### Western blot

Snap-frozen ventricular tissue was homogenized in RIPA buffer (50 mM Tris pH 7.4, 150 mM NaCl, 1 mM EDTA, 1 % Triton X-100, 1 % sodium deoxycholate, 2 mM NaF, 200 µM Na_3_VO_4_, 0.1% sodium dodecyl sulfate, 5 mM *_N_*-ethylmaleimide) supplemented with HALT protease and phosphatase inhibitor cocktail (Thermo Fisher) at a concentration of 100 mg of tissue per ml of buffer. Following sonication, lysates were clarified via centrifugation at 10,000 x g for 20 min at 4 °C, and DC protein assay (Bio-rad) was used to measure protein concentration. 4X Bolt LDS sample buffer (Thermo Fisher) with 400 mM DTT (Sigma Aldrich) was added to lysates, which were then heated at 70 °C for 10 min. Samples underwent SDS-PAGE with NuPAGE Bis-Tris Midi 4-12 % gradient gels and MES running buffer (Thermo Fisher). Proteins were then transferred to LF-PVDF membranes (Bio-rad), with membranes methanol-fixed and air-dried post-transfer. Following methanol reactivation, membranes were blocked for 1 h at room temperature in 5 % nonfat milk (Carnation) in TNT buffer (0.1 % Tween 20, 150 mM NaCl, 50 mM Tris pH 8.0). Primary antibody incubation was conducted overnight at 4 °C with the following antibodies: rabbit anti-Cx43 (1:4,000; Sigma Aldrich), rabbit anti-HIF-2α (1:1,000; Cell Signaling Technology), rabbit anti-VHL (1:1,000; Thermo Fisher), and mouse anti-GAPDH (1:1,000; Proteintech). Following primary antibody incubation, membranes were rinsed twice and washed 3 × 10 min in TNT buffer. Secondary antibody incubation was conducted for 1 h at room temperature with antibodies conjugated to AlexaFluor555 or AlexaFluor647 (1:5,000; Thermo Fisher). Washes were conducted in TNT buffer as previously described. Images were acquired using a Chemidoc MP imaging system (Bio-rad).

### RT-qPCR

RNA was isolated as above for RNA sequencing. cDNA was generated with iScript Advanced cDNA synthesis kit for RT-qPCR (Bio-Rad) according to manufacturer’s instructions. Real-time PCR was performed with Luna Universal qPCR Master Mix (New England Biolabs) on a QuantStudio 6 Flex System (Thermo Fisher) employing the following primers: *Cacna1c*, Mm.PT.58.8608981 IDT; *Scn5a*, Mm.PT.58.30418114 IDT; *Cdh2*, Mm.PT.58.12378183 IDT; *Pkp2*, Mm.PT.58.11914062 IDT; T*jp1*, Mm.PT.58.12952721 IDT; *Gja1*, Mm.PT.58.5955325 IDT; *Gapdh*, Mm.PT.39a.1 IDT; *Kcnq1*, Mm.PT.58.9135325 IDT; *Myh6* Fwd CCA ACA CCA ACC TGT CCA AGT, *Myh6* Rev AGA GGT TAT TCC TCG TCG TGC AT; *Myh7* Fwd CTC AAG CTG CTC AGC AAT CTA TTT, Myh7 Rev GGA GCG CAA GTT TGT CAT AAG T; *Nppa* Fwd TTC CTC GTC TTG GCC TTT TG, *Nppa* Rev CCT CAT CTT CTA CCG GCA TC; *Nppb* Fwd GTC CAG CAG AGA CCT CAA AA, *Nppb* Rev AGG CAG AGT CAG AAA CTG GA.

